# Radiation synergizes with BET inhibition to stimulate durable, systemic anti-tumor immunity in murine cancer models

**DOI:** 10.64898/2026.02.16.706212

**Authors:** Nicole McCuen, Chantal Vidal, Kamal Pandey, SM Nashir Udden, Yu-Lun Liu, Prasanna G. Alluri

**Author notes:** Address correspondence to: Prasanna G. Alluri, MD, PhD. The authors have declared that no conflict of interest exists.

## Abstract

Most patients with breast cancer (BC) and soft tissue sarcoma (STS) harbor immunologically cold tumors and do not respond to existing immunotherapies such as immune checkpoint inhibitors (ICIs) as a monotherapy. Consequently, prolonged treatment with highly toxic multiagent chemotherapy, with or without ICIs, remains the mainstay of systemic therapy in such patients. Therefore, there is an acute clinical need for novel chemotherapy-free immunotherapy regimens with high efficacy and minimal toxicity. Here, employing an *in vivo* drug screen, we identify that a short course of radiation therapy (RT) synergizes with pharmacological bromodomain and extraterminal (BET) inhibition to elicit a strong systemic anti-tumor immunity and long-term immunological memory in a CD8+ T cell-dependent manner in murine models of both BC and STS. Mechanistic studies reveal that RT + BET inhibition accentuates RT-induced DNA damage and micronuclei formation, increases Major Histocompatibility Complex class I and II expression on macrophages, enhances translocation of calreticulin to the plasma membrane, and blocks RT-induced Programmed Death-Ligand 1 (PD-L1) overexpression on tumor cells, thereby promoting immunogenic cell death. Our data suggest that a combination of RT + BET inhibition promotes robust anti-tumor immunity and immunological memory in immunologically cold tumors, thereby opening potential avenues for clinical translation.

## Introduction

Immunotherapies have revolutionized the care of many cancer patients. For example, immune checkpoint inhibitors (ICIs) significantly improve overall survival in advanced melanoma patients (1). In tumors exhibiting microsatellite instability, ICI regimens have proven more effective than chemotherapy regimens (2, 3). However, in poorly immunogenic cancers such as breast cancer (BC) and soft tissue sarcomas (STS), ICIs fail to afford clinical benefit in most patients (4, 5). Even in a small subset of BC patients such as those with Triple Negative Breast Cancer (TNBC) in whom immunotherapies are Food and Drug Administration (FDA)-approved, ICIs require co-administration with a multiagent chemotherapy backbone to improve clinical outcomes. For example, in patients with localized TNBC, the existing standard-of-care (SOC) KEYNOTE-522 regimen involves co-administration of pembrolizumab, an anti-PD-1 antibody, with a multiagent cytotoxic chemotherapy regimen comprised of paclitaxel, carboplatin, doxorubicin or epirubicin and cyclophosphamide in the preoperative setting (6). While addition of pembrolizumab to chemotherapy improved 5-year overall survival from 81.7% to 86.6%, a significant subset of patients fail to achieve a pathological complete response (pCR) and face a high risk of recurrence (6). Additionally, this current SOC chemo-immunotherapy regimen comes at a very high toxicity penalty to patients, with over 80% of patients experiencing Grade ≥ 3 toxicity. Over 90% of the observed toxicity in these patients is attributable to chemotherapy (6). Similarly, most patients with STS harbor immunologically cold tumors and, with the exception of rare subtypes such as Alveolar Soft Part Sarcoma (ASPS) (7), immunotherapies such as ICIs fail to provide clinical benefit (5, 8). Therefore, highly toxic chemotherapy regimens remain the mainstay of systemic therapy in such patients (9). In a systemic review of clinical trials of patients with STS, Grade ≥ 3 toxicity rate was 58% for anthracycline-based chemotherapy regimens and 84% for kinase inhibitor regimens (10). Therefore, there is an urgent clinical need to develop novel chemotherapy-free immunotherapy regimens that elicit a strong anti-tumor immune response and long-term immunological memory with minimal toxicity. The ideal characteristics of such a novel immunotherapy regimen include the ability to stimulate a strong systemic anti-tumor immune response in otherwise immunologically cold tumors without the need for a chemotherapy backbone, short total treatment duration (which increases patient convenience and compliance), a non-invasive route of administration (which obviate the need for intravenous infusions or intratumoral injections), a durable and long-term immunological memory (which reduces the risk of recurrence), and minimal toxicity.

Radiation therapy (RT) is commonly administered to eliminate residual microscopic disease after surgery in patients with main types of cancer such as BC and STS (11, 12). However, when targeted against an intact tumor, RT also has the potential to mediate a systemic anti-tumor immune response through multiple mechanisms (13), including the induction of immune cell death (14), release of double-strand DNA (15), release of danger-associated molecular patterns (DAMPs) (16), stimulation of dendritic cell maturation (17), and cross-priming of cytotoxic T lymphocytes (CTLs) (18). Thus, RT delivered against a local tumor has the potential to promote elimination of tumors outside the irradiated field. For instance, a combination of irradiation and ICI synergistically promote antitumor immunity in mice (19) as well as in patients (20–22) although such occurrences are exceedingly rare, likely due to competing immune-stimulatory and immunosuppressive effects of RT on the tumor microenvironment (23, 24).

Here, employing an *in vivo* drug screen, we identify that a short course of pharmacological bromodomain and extra-terminal domain (BET) inhibition synergizes with tumor-directed RT to stimulate a robust anti-tumor immune response and long-term immunological memory in a CD8+ T cell dependent fashion in multiple immunologically cold murine models, including both BC and STS. Furthermore, we show that BET inhibition enhances immune-stimulatory effects of RT while blunting it’s immunosuppressive effects, thereby positively influencing several steps involved in the immune activation cycle.

## Results

### Pharmacological BET inhibition synergizes with tumor-directed RT to inhibit primary tumor growth in multiple syngeneic mouse models

Epigenetic changes regulate many key steps involved in the generation of anti-tumor immunity (25), including but not limited to CD4+ T cell differentiation (26), CD8+ T cell differentiation (27), dedifferentiation of cytotoxic T cells into memory T cells (28), T cell exhaustion (29), myeloid cell differentiation (30), and macrophage polarization (31). Therefore, there is a strong interest in exploiting epigenetic therapeutics to promote anti-tumor immunity (32). Since RT has both immune-stimulatory (13) and immunosuppressive effects (23), we hypothesized that combining RT with an epigenetic modulator which enhances its immune-stimulatory effects while blocking the immunosuppressive effects will promote a robust anti-tumor immune response and long-term immunological memory in otherwise immunologically cold tumors.

To identify potential epigenetic modulators that synergize with RT to elicit a potent anti-tumor immune response, we carried out an *in vivo* drug screen against 4T1, a highly aggressive syngeneic TNBC mouse model, using a panel of 19 epigenetic inhibitory drugs with diverse targets (Figure 1A and B). Since our goal was to identify a treatment regimen that only requires few administrations to stimulate an anti-tumor immune response (as opposed to existing treatment regimens that involve prolonged treatment), our study was designed to administer a sub-therapeutic radiation dose of 8 Gy x 2 to orthotopic 4T1 tumors. Similarly, since our goal is to identify drugs that synergize with RT to induce an anti-tumor immune response (as opposed to relying on systemic drug effects), the drug administration was limited to 4 days, with the treatment starting a day before the start of RT and ending on the last day of RT. Thus, RT was administered on days 2 and 4 in combination with a library drug on days 1-4. No treatment was administered to mice after day 4 (Figure 1A). The study endpoint was tumor volume two weeks after treatment.

**Figure 1:**
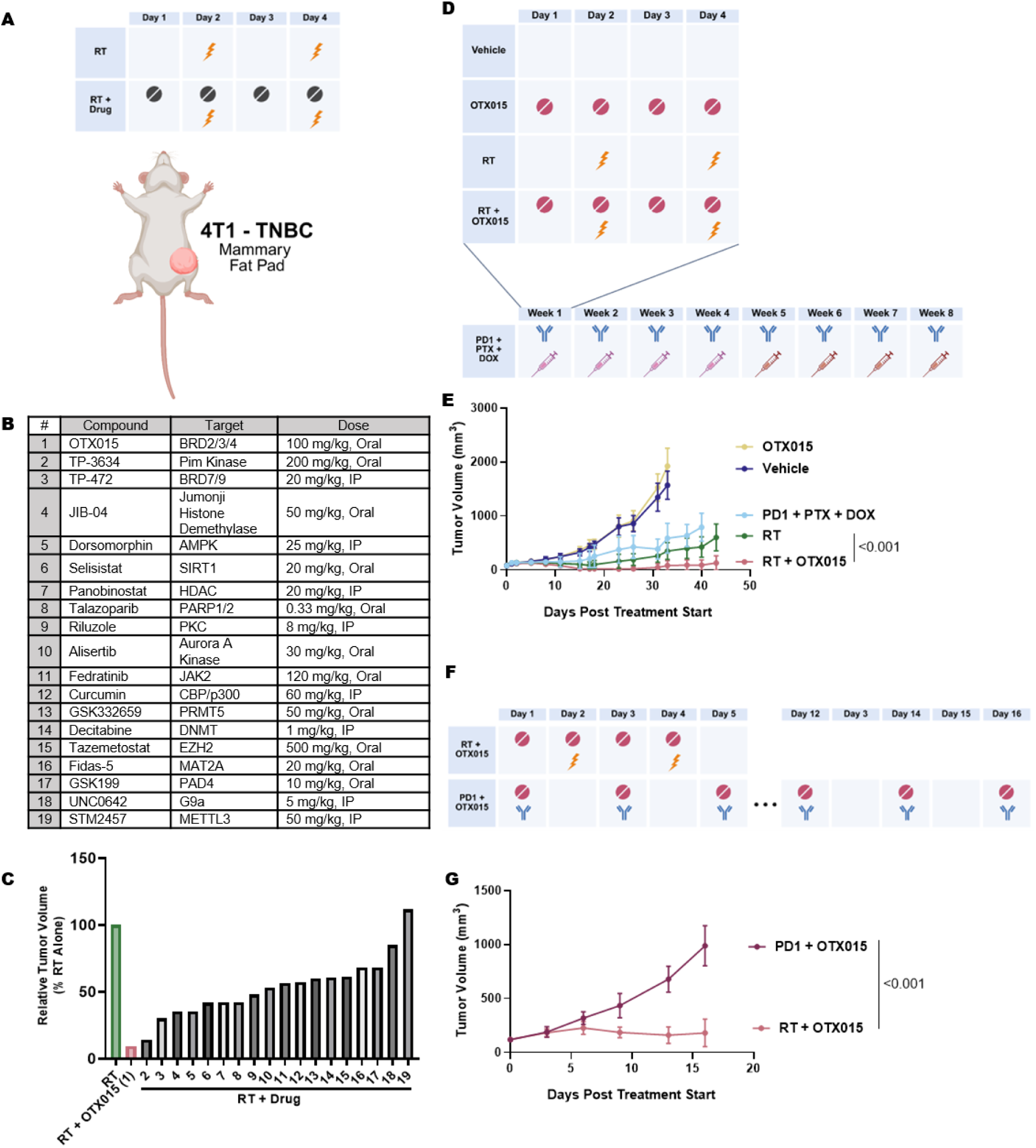
RT synergizes with OTX015 to inhibit the growth of 4T1 tumors. (A): Schematic of experimental design. Female BALB/c mice bearing orthotopic 4T1 tumors in the mammary fat pad were randomized when tumors reached 100-150 mm^3^ and treated with RT alone (8 Gy on days 2 and 4) or RT (8 Gy on days 2 and 4) in combination with a library drug (drug treatment on days 1-4) as indicated (n=1 mouse per treatment). (B): A list of library drugs used in the screen along with their targets. (C): Two weeks after treatment, tumor volumes were measured and compared to RT alone treatment as reference. The number on the x-axis of the bar graph corresponds to the drug number in the table shown in B. (D): Schematic of experimental design. Female, BALB/c mice bearing orthotopic 4T1 tumors were randomized when tumors reached 100-150 mm^3^ to the following treatment arms: Vehicle, OTX015 (100 mg/kg by oral gavage on days 1-4), RT (8 Gy on days 2 and 4), RT + OTX015, or an 8-week chemo-immunotherapy regimen (anti-PD1 antibody, 200 µg/dose i.p. weekly for 8 weeks; Paclitaxel, or PTX, 10 mg/kg IP weekly during weeks 1-4; Doxorubicin, or DOX, 2 mg/kg i.p. weekly during weeks 5-8), as indicated. (E): The results are plotted as average tumor volume for each treatment arm ± SD (n=8 mice/arm). Tumor growth curves were analyzed using linear mixed-effects models to account for repeated measurements within each mouse, with treatment and time as fixed effects and mouse as a random effect. Pre-specified pairwise comparisons were performed using estimated marginal means with Benjamini–Hochberg correction applied within each comparison family. (F): Female, BALB/c mice bearing orthotopic 4T1 tumors were randomized when tumors reached 100-150 mm^3^ and treated with anti-PD1 antibody (200 µg/dose, 3x/week) + OTX015 (100mg/kg by oral gavage, 3x/week) or RT (8 Gy on days 2 and 4) + OTX015 (100 mg/kg by oral gavage on days 1-4). (G): The results are plotted as average tumor volume for each treatment arm ± SD (n=8 mice/arm). The same statistical analysis described in E was used. Abbreviations: BRD – bromodomain and extraterminal domain; AMPK – 5’ adenosine monophosphate-activated protein kinase; SIRT1 – sirtuin 1; HDAC – histone deacetylase; PARP – poly (ADP-ribose) polymerase; PKC – protein kinase C; JAK2 – Janus kinase 2; CBP – cAMP response element-binding protein (CREB)-binding protein; PRMT5 – protein arginine methyltransferase 5; DNMT – DNA methyltransferase; EZH2 – enhancer of zeste homolog 2; MAT2A – methionine adenosyltransferase 2A; PAD4 – protein arginine deaminase 4; METTL3 – N(6)-adenosine-methyltransferase; i.p. – intraperitoneal injection

In this screen, OTX015, a small molecule inhibitor of bromodomain and extrateminal (BET) family of proteins (33), emerged as the top hit. The combination of RT + OTX015 caused an approximately 90% reduction in tumor volume relative to RT alone, despite only four total doses of OTX015 administration (Figure 1C). The BET family proteins such as BRD4 are epigenetic regulators that bind to acetylated histones and recruit transcription factors and elongation complexes to promote the expression of many oncogenic drivers in a context dependent fashion (34). In addition, BRD4 plays an important role in DNA damage response and is essential for repair of double strands breaks through non-homologous end joining repair (NHEJ) pathway (35). BET proteins also play a key role in attenuating immune surveillance against tumors (36) and in promoting an immunosuppressive tumor microenvironment (TME) (37–39). Despite their early promise, BET inhibitors have failed to achieve clinic success as a single agent, both due to poor efficacy and high toxicity (40). Therefore, current efforts are focused on identification of novel BET inhibitor combination regimens that improve efficacy and have led to promising results in Phase 2 and Phase 3 clinical trials (41, 42).

We validated the findings from this drug screen in a comprehensive study with all the appropriate controls. 4T1 tumors were implanted in the mammary fat pad of female BALB/c mice. When tumors reached 100-150 mm^3^, mice were randomized to the following treatment arms: Vehicle, OTX015 alone (100 mg/kg on days 1-4), tumor-directed RT alone (8 Gy on days 2 and 4), chemo-immunotherapy [8-week regimen of anti-PD1 antibody (PD1), paclitaxel (PTX) and doxorubicin (DOX)] or RT + OTX015 (Figure 1D). Consistent with our findings from the initial screen, RT + OTX015 induced a potent tumor growth suppression and outperformed all other arms, including an 8-week regimen of chemo-immunotherapy (Figure 1E). In a separate study, we also compared the RT + OTX015 regimen to anti-PD1 + OTX015 regimen since previous preclinical studies have shown pharmacological BET inhibition synergizes with PD-1 blockade to inhibit leukemia progression (43). Again, we found that RT + OTX015 treatment over 4 days exhibited far superior tumor control relative to continuous administration of anti-PD1 antibody + OTX015 treatment (Figure 1F and G). These findings were further validated in a second BC model, an Estrogen Receptorα-positive (ER+) syngeneic mouse model, MXT+, implanted orthotopically in female BDF1 mice (44). In this model, although RT alone and OTX015 alone exerted modest tumor control, RT + OTX015 completely inhibited tumor growth (Figure S1). These findings strongly suggest that pharmacological BET inhibition synergizes with RT to inhibit the growth of both ER+ and Triple Negative breast tumors.

To assess the generalizability of our findings and establish whether the combination of RT + OTX015 inhibits tumor growth across diverse tumor types, we evaluated the efficacy of this combination in an additional syngeneic mouse model of TNBC (EMT6) and STS (MCA205), thereby capturing the heterogeneity of cancers originating from both epithelial cells and mesenchymal cells, respectively. While the EMT6 model received the aforementioned 4-day regimen, to account for the radioresistant nature of STS, we employed a 6-day combination regimen for the MCA205 model comprised of three doses of radiation (8 Gy on days 2,4 and 6) and six total doses of OTX015 (on days 1-6) (Figure 2A and B). In both EMT6 and MCA205 models, the combination of RT + OTX015 completely inhibited tumor growth, although neither RT alone or OTX015 alone achieved tumor growth arrest (Figure 2C and D). At ∼ 7 weeks after treatment, 10/10 MCA205-bearing mice treated with RT + OTX015 were alive while only 2/10 mice receiving RT and 0/10 mice receiving OTX015 were alive (Figure S2). Remarkably, RT + OTX015 treatment completely eradicated tumors in 2/10 mice bearing EMT6 tumors and 4/10 mice bearing MCA205 tumors. Furthermore, mice receiving RT + OTX015 did not show any signs of toxicity such as body weight loss, complete blood counts, or evidence of serum biochemistry abnormalities that reflect bone marrow, liver or kidney toxicities (Figure S3A-C).

**Figure 2:**
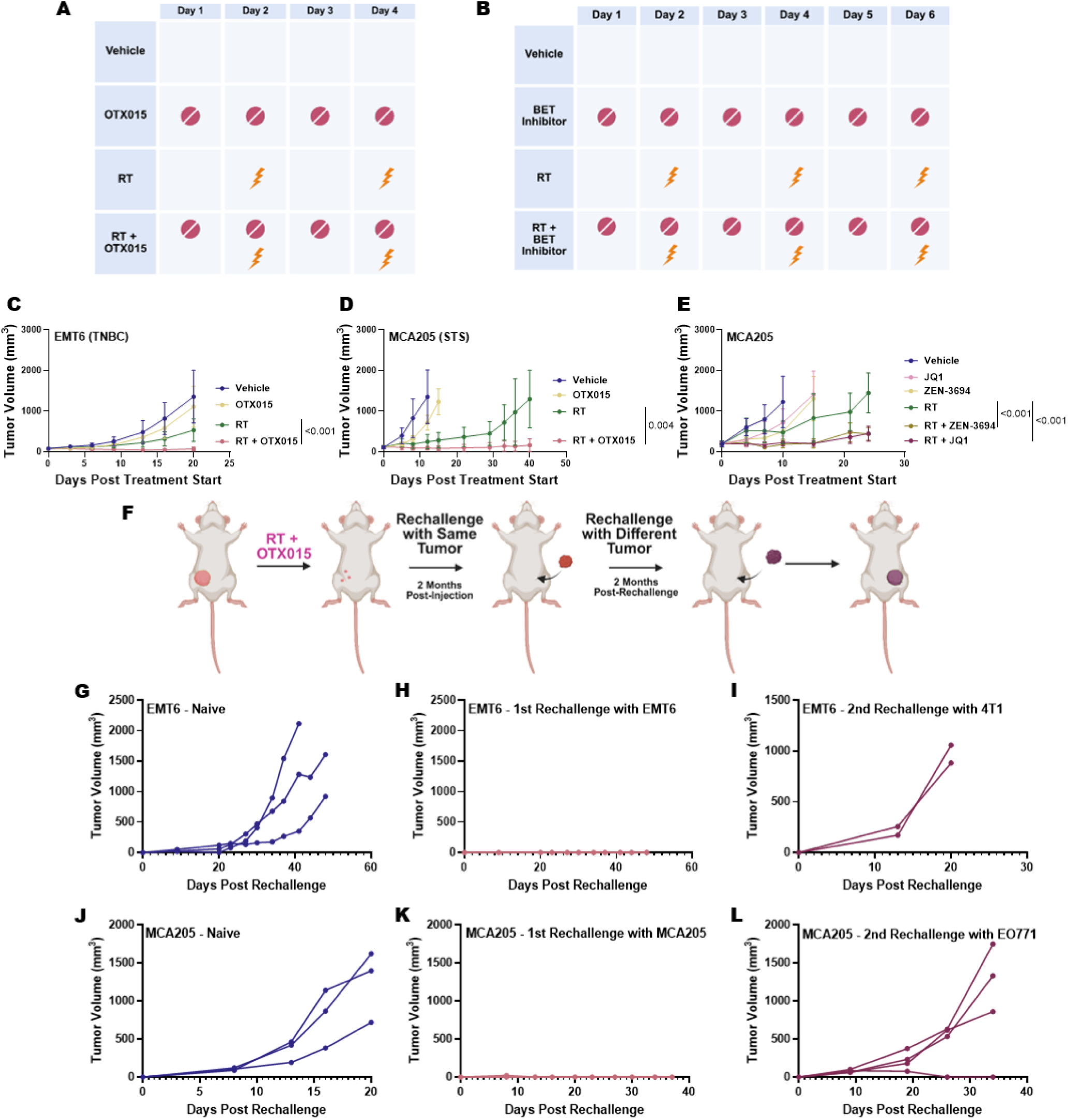
RT synergizes with BET inhibitors and promotes durable anti-tumor immune response across diverse tumor types. (A-B): Schematic depictions of a 4-day (A) or 6-day (B) treatment regimen in which 8 Gy RT was administered on days 2 and 4 (A) or days 2, 4 and 6 (B) and the BET inhibitor administered daily on days 1-4 (A) or 1-6 (B), as indicated. (C): Female, BALB/c mice bearing orthotopic EMT6 tumors were randomized when tumors reached 100-150 mm^3^ to the following treatment arms: Vehicle, RT (8 Gy on days 2 and 4), OTX015 (100 mg/kg by oral gavage on days 1-4) or RT + OTX015. The results are plotted as average tumor volume for each treatment arm ± SD (n=8-10 mice/arm) where 2/10 mice were cured by RT + OTX. (D): Female, C57BL6 mice bearing subcutaneous MCA205 tumors in the right flank were randomized when tumors reached 100-150 mm^3^ to the following treatment arms: Vehicle, RT (8 Gy on days 2, 4 and 6), OTX015 (100 mg/kg by oral gavage on days 1-6), or RT + OTX015. The results are plotted as average tumor volume for each treatment arm ± SD (n=8-10 mice/arm) where 4/10 mice were cured by RT + OTX015. (E): Female, C57BL6 mice bearing subcutaneous MCA205 tumors in the right flank were randomized when tumors reached 100-150 mm^3^ to one of the following treatment arms: Vehicle, RT (8 Gy on days 2, 4 and 6) ZEN-3694 (100 mg/kg by oral gavage on days 1-6), JQ1 (50 mg/kg IP on days 1-6), RT+ ZEN-3694, or RT + JQ1. The results are plotted as average tumor volume for each treatment arm ± SD (n=7 mice/arm). For C, D and E, tumor growth curves were analyzed using linear mixed-effects models to account for repeated measurements within each mouse, with treatment and time as fixed effects and mouse as a random effect. Pre-specified pairwise comparisons were performed using estimated marginal means with Benjamini–Hochberg correction applied within each comparison family. (F): Schematic depiction of tumor rechallenge experiment. (G): Naive female, BALB/c mice (n=3) were injected orthotopically with EMT6 cells. (H): Two months after initial tumor injection, female BALB/c mice cured of EMT6 tumors (from C) following treatment with RT + OTX015 (n=2), were rechallenged with new EMT6 cells in the contralateral mammary fat pad. (I): Two months following rechallenge in H, the same female, BALB/c mice which resisted EMT6 rechallenge were injected with 4T1 cells. (J): Naive female, C57BL6 mice (n=3) were injected subcutaneously with MCA205 cells. (K): Two months after initial tumor injection, female C57BL6 mice cured of MCA205 tumors (from D) following treatment with RT + OTX015 (n=4), were rechallenged with new MCA205 cells in the contralateral flank. (L): Two months following rechallenge in K, the same female, C57BL6 mice which resisted MCA205 rechallenge were injected with E0771 cells.

To assess whether the efficacy of RT + OTX015 combination is generalizable to BET inhibitors as a class, we evaluated the efficacy of two additional BET inhibitors, JQ1 (a probe compound) and ZEN-3694 (a clinical grade BET inhibitor), in combination with RT in MCA205-bearing mice. ZEN-3694 is currently in active clinical development and has advanced to Phase 2 clinical trials for treatment for many types of cancers, including NUT carcinoma (NCT05019716), advanced prostate cancer (NCT04986423), lung cancer (NCT05607108) and ovarian cancer (NCT05071937). Similar to OTX015, both RT + JQ1 and RT + ZEN-3694 achieved superior tumor control relative to RT alone and drug alone. Thus, although 0% of mice in the vehicle, JQ1 and ZEN-3694 arms, and only 43% of mice in the RT arm were alive at the time of study termination, 100% of mice in both the RT + JQ1 and RT + ZEN-3694 arms were alive. These findings suggest that the efficacy of the combination treatment is related to pharmacological BET inhibition and not specific to OTX015 (Figure 2E, Figure S4).

### RT + OTX015 elicits robust systemic anti-tumor immunity and long-term immunological memory

To test whether RT + OTX015 elicits a systemic anti-tumor immune response in addition to enhanced local control of the treated primary tumors, we performed tumor rechallenge experiments in mice cured of EMT6 and MCA205 tumors following treatment with RT + OTX015. Two months following initial tumor injection, mice cured by RT + OTX015 were re-injected with the same cancer cell line in the contralateral mammary fat pad (EMT6) or flank (MCA205) (Figure 2F). Simultaneously, as a positive control, treatment naïve mice were injected with the same tumor cells (Figure 2G and J). Across both models, 100% of cured mice resisted rechallenge and failed to regrow tumors while 100% of treatment naïve mice rapidly developed tumors (Figure 2H and K).

To test the specificity of resistance to rechallenge, mice that were initially cured of and resisted rechallenge with EMT6 cells were subjected to a second rechallenge two months later with 4T1 cells, which are also syngeneic to BALB/c mice. 2/2 mice injected with 4T1 cells rapidly grew tumors although they had previously resisted EMT6 rechallenge (Figure 2I). Similarly, mice that were initially cured of and resisted rechallenge with MCA205 were rechallenged with E0771 cells, which are also syngeneic to C57BL/6 mice. 3/4 mice rechallenged with E0771 cells rapidly developed tumors (Figure 2L). These findings strongly suggest that RT + OTX015 induces long term immunological memory in a largely tumor-specific manner.

Next, to establish whether RT + OTX015 treatment stimulates a robust systemic anti-tumor immune response which controls distant non-irradiated tumors, we implanted MCA205 tumors bilaterally in female C57BL/6 mice and treated them with RT to the right flank alone, along with concurrent oral administration of vehicle or OTX015 (Figure 3A). As expected, treatment with RT + OTX015 induced a more potent tumor growth suppression relative to RT + vehicle (Mean tumor weight of 208 mg with RT vs 94 mg with RT + OTX015) (Figure 3B). Remarkably, the unirradiated tumors in mice treated with RT + OTX015 showed significant tumor growth suppression relative to mice receiving RT + vehicle (mean tumor weight of 819 mg in mice treated with RT + vehicle vs 329 mg in mice receiving RT + OTX015) (Figure 3C), although treatment with OTX015 alone failed to achieve significant tumor growth suppression in all our studies. These findings suggest that RT + OTX015 elicits a systemic anti-tumor immune response and controls both the irradiated local tumors as well as non-irradiated distant tumors. Furthermore, we found a significant increase in both CD8+ and CD4+ T cells in tumors treated with RT + OTX015 relative to RT alone (Figure 3D-F). Remarkably, mice treated with RT + OTX015 also showed evidence of increased infiltration of both CD8+ and CD4+ T cells in the non-irradiated tumors. (Figure 3D, G and H).

**Figure 3:**
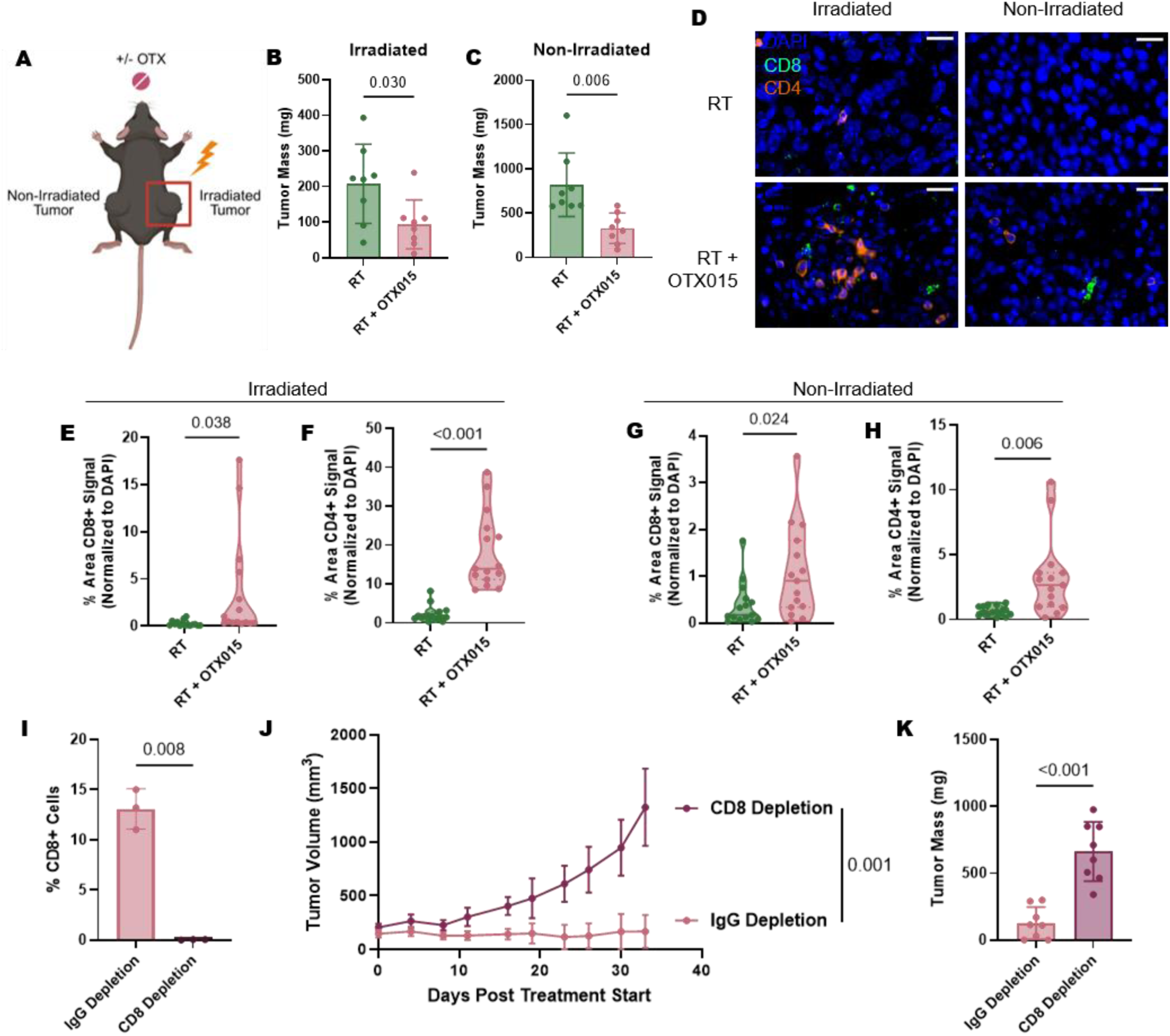
RT + OTX015 elicits a systemic anti-tumor immune response in a CD8+ T cell-dependent manner. (A): Female C57BL6 mice were injected with MCA205 cells in both flanks. When tumors reached 100-150 mm^3^, mice were randomized for treatment with RT (8 Gy on days 2, 4, and 6 to right flank alone) or RT + OTX015 (RT= 8 Gy on days 2, 4 and 6 to right flank alone; OTX015: 100 mg/kg by oral gavage on days 1-6). (B-C): Two weeks after treatment, all tumors were excised and weighed (n=8 mice/arm). (D): Tumors were then stained for DAPI, CD8, and CD4 and imaged using immunofluorescent microscopy, and representative images are shown. Scale bar is 20 microns. (E-H): Immune infiltration was quantified by calculating the % area of fluorescence intensity normalized to DAPI for each marker (n=15 images for each arm, 3 tumors were stained and 5 images per tumor were quantified to account for variation between biological replicates and spatial distribution of immune cells). Infiltration of CD8+ T cells (E) and CD4+ T cells (F) of irradiated tumors and CD8+ T cells (G) and CD4+ T cells (H) of non-irradiated T cells are plotted. (I): Female, C57BL6 mice were administered either IgG or CD8 depletion antibody (200 µg/dose) a day before tumor injection, the day of tumor injection, and twice weekly thereafter. Depletion was confirmed using flow cytometry staining of peripheral blood from a sampling of mice (n=3 mice/arm) for CD8. (J): IgG or CD8 antibody-treated mice were treated with the 6-day regimen of RT + OTX015 as described previously. The results are plotted as average tumor volume for each treatment arm ± SD (n=8 mice/arm). (K): Tumor mass at treatment endpoint. For Figure 3B-I, P-values were determined using an unpaired, 2-tailed *t*-test with Welch’s correction. For Figure 3J, P-value was determined using linear mixed-effects models.

Based on these observations, we hypothesized that the anti-tumor response mediated by RT + OTX015 is driven by adaptive immunity. To definitively establish the functional contribution of CD8+ T cells in mediating the anti-tumor activity of RT + OTX015, we performed an *in vivo* immune cell depletion study. Depletion of CD8+ T cells was confirmed by flow cytometry analysis of peripheral blood (Figure 3I). As expected, mice administered with IgG (control) antibody showed potent tumor growth inhibition following treatment with RT + OTX015. However, the tumor control was abrogated in mice depleted of CD8+ T cells (Figure 3J and K). These data definitively establish that the anti-tumor activity of RT + OTX015 is dependent on CD8+ T cell function.

### OTX015 enhances RT-induced DNA damage and micronuclei formation

DNA damage is a potent activator of innate immune response. RT-induced DNA damage and micronuclei release double stranded DNA into the cytoplasm and stimulate expression of Type I interferons and inflammatory cytokines (45). Suppression of DNA repair following RT-induced DNA damage also has the potential to increase tumor mutational burden and generation of neoantigens, which could promote tumor-specific immunity (46, 47). Given the essential role of BET proteins such as BRD4 in the repair of DNA double strand breaks through non-homologous end joining (NHEJ) repair pathway (35), we hypothesized that the immune effects of RT + OTX015 are mediated, at least in part, through accentuation of RT-induced DNA damage. Consistent with this hypothesis, OTX015 treatment significantly increased RT-induced γH2AX foci in MCA205 cells both at 30 minutes and 120 minutes following RT, suggesting persistence of unrepaired DNA damage in these cells (Figure 4A and B). Next, we explored the impact of RT + OTX015 on micronuclei formation in MCA205 cells. Micronuclei develop from genotoxic stress-induced chromatin fragmentation (45). Consistent with our findings that OTX015 accentuates RT-induced DNA damage, MCA205 cells treated with RT + OTX015 showed higher number of micronuclei relative to both RT alone and OTX015 alone (Figure 4C and D).

**Figure 4:**
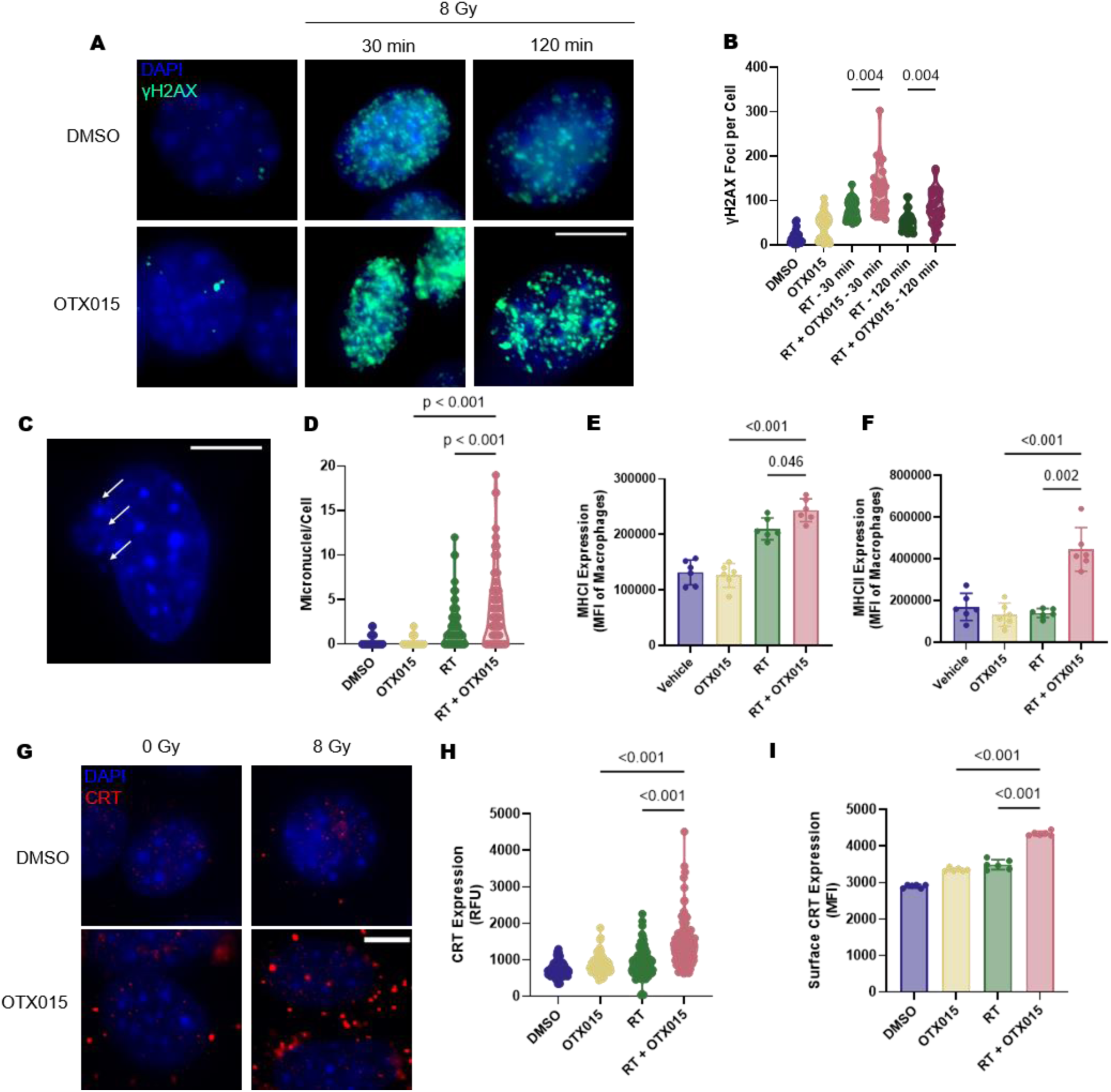
RT + OTX015 promotes immunological cell death. (A-B): MCA205 cells were treated with DMSO, OTX015 (0.5 µM), RT (8 Gy), or RT + OTX015. At 30 minutes and 120 minutes post-treatment, cells were fixed and stained with DAPI and γH2AX antibodies and imaged using immunofluorescent microscopy. Scale bar is 10 microns. γH2AX foci (n=25) were counted and plotted as foci per cell. (C-D): MCA205 cells were treated with DMSO, OTX015, RT or RT + OTX015 as described above. 5 days post-RT, cells were fixed and stained with DAPI. Micronuclei were quantified using immunofluorescence microscopy and plotted as number of micronuclei per cell. (E-F): Female, C57BL6 mice (n=6) bearing MCA205 tumors were randomized and treated with Vehicle, OTX015 (100 mg/kg by oral gavage on days 1-6), RT (8 Gy on days 2, 4 and 6), or RT + OTX015. Two weeks after treatment, tumors were harvested, digested, and analyzed by flow cytometry for MHCI (E) and MHCII (F) expression on macrophages as defined by CD45+CD11b+F480+ cells. (G-I): MCA205 cells were treated with DMSO, RT, OTX015 or RT+OTX015 as described in A. 24 hours post-treatment, cells were fixed and stained with DAPI and calreticulin (CRT) and imaged using immunofluorescence microscopy. Scale bar is 10 microns. The relative fluorescence intensity of CRT per cell (n=100) was quantified with ImageJ and plotted. (H): MCA205 cells (n=6 wells per treatment) under the same conditions were collected 24 hours post-treatment and analyzed by flow cytometry for surface expression of CRT. The results are plotted as MFI per biological replicate (I). Overall differences among groups were assessed using one-way ANOVA. Prespecified pairwise comparisons were performed using unpaired Welch’s *t*-tests to account for unequal variances. Abbreviations: MFI – Mean Fluorescence Intensity

### RT + OTX015 stimulates MHC class I and II expression on macrophages and promotes immunogenic cell death

Next, we evaluated the impact of RT + OTX015 on expression of MHC class I and II expression on antigen presenting cells. Remarkably, mice bearing MCA205 tumors showed increased MHC class I and II expression on macrophages following treatment with RT + OTX015 relative to both RT alone and OTX015 alone (Figure 4E and F). Next, we asked if the enhanced DNA damage and increased MHC class I and II expression induced by RT + OTX015 promotes immunogenic cell death. Immunogenic cell death results from initiation of an immune response against the damaged cells through release of damage associated molecular patterns (DAMPs) and antigen presentation (14). One of the mechanisms by which this occurs is through translocation of the calreticulin (CRT) to the plasma membrane (48). CRT serves as a chaperone in the endoplasmic reticulum (ER), but following ER stress, it translocates to the plasma membrane to serve as a damage-associated molecular pattern (DAMP), aid in MHCI folding, and recruitment of antigen-presenting cells (49). 24 hours following treatment, we observed enhanced CRT accumulation by immunofluorescence imaging in MCA205 cells treated with RT + OTX015 relative to both RT alone and OTX015 alone (Figure 4G and H). We confirmed that the accumulation of CRT is occurring at the plasma membrane rather than intracellularly through flow cytometry analysis of cell surface markers, which confirmed a significant increase is CRT surface expression in cells treated with RT + OTX015 (Figure 4I). These data strongly suggest that RT + OTX015 induces immunogenic cell death by increasing MHC class I and II expression and promoting CRT translocation to the plasma membrane, thereby enhancing antigen presentation to T cells.

### OTX015 blocks RT-induced PD-L1 upregulation

Even with adequate antigen presentation and T cell activation, cancer cells may evade immune recognition through upregulation of immune checkpoints such as PD-L1 (50). While RT may promote several steps involved in immune activation, it also has the potential to promote immunosuppression, especially through upregulation of PD-L1 expression on tumor cells (19). We observed a dramatic time-dependent upregulation of PD-L1 expression on tumor cells following RT across multiple cancer cell lines, including both mouse and human cell lines (Figure 5A-C), which is consistent with previous reports (51). However, concurrent treatment of cells with RT and OTX015 abrogated RT-induced PD-L1 overexpression (Figure 5D-F). Next, we investigated whether RT-induced PD-L1 overexpression is mediated by BRD4. Using a chromatin immunoprecipitation (ChIP) assay, we observed a significant increase in BRD4 recruitment to the PD-L1 promoter following RT in human TNBC cell line MDA-MB-231 (Figure 5G and H). Concurrent OTX015 administration abrogated this RT-induced BRD4 recruitment (Figure 5H). While previous studies have shown that BRD4 inhibition suppresses PD-L1 expression (52, 53), to our knowledge, our study is the first to demonstrate that RT-induced PD-L1 overexpression is BRD4 dependent. Thus, we uncover a novel mechanism for RT-induced PD-L1 upregulation, which is targetable through BET inhibition. Overall, our findings highlight the dual role of BET inhibition in both accentuating RT-induced immune activation (through enhanced DNA damage and micronuclei formation, stimulation of MHC I and II expression, and promotion of calreticulin translocation to the nucleus) and dampening of RT-induced immunosuppression (through abrogation of RT-induced PD-L1 overexpression).

**Figure 5:**
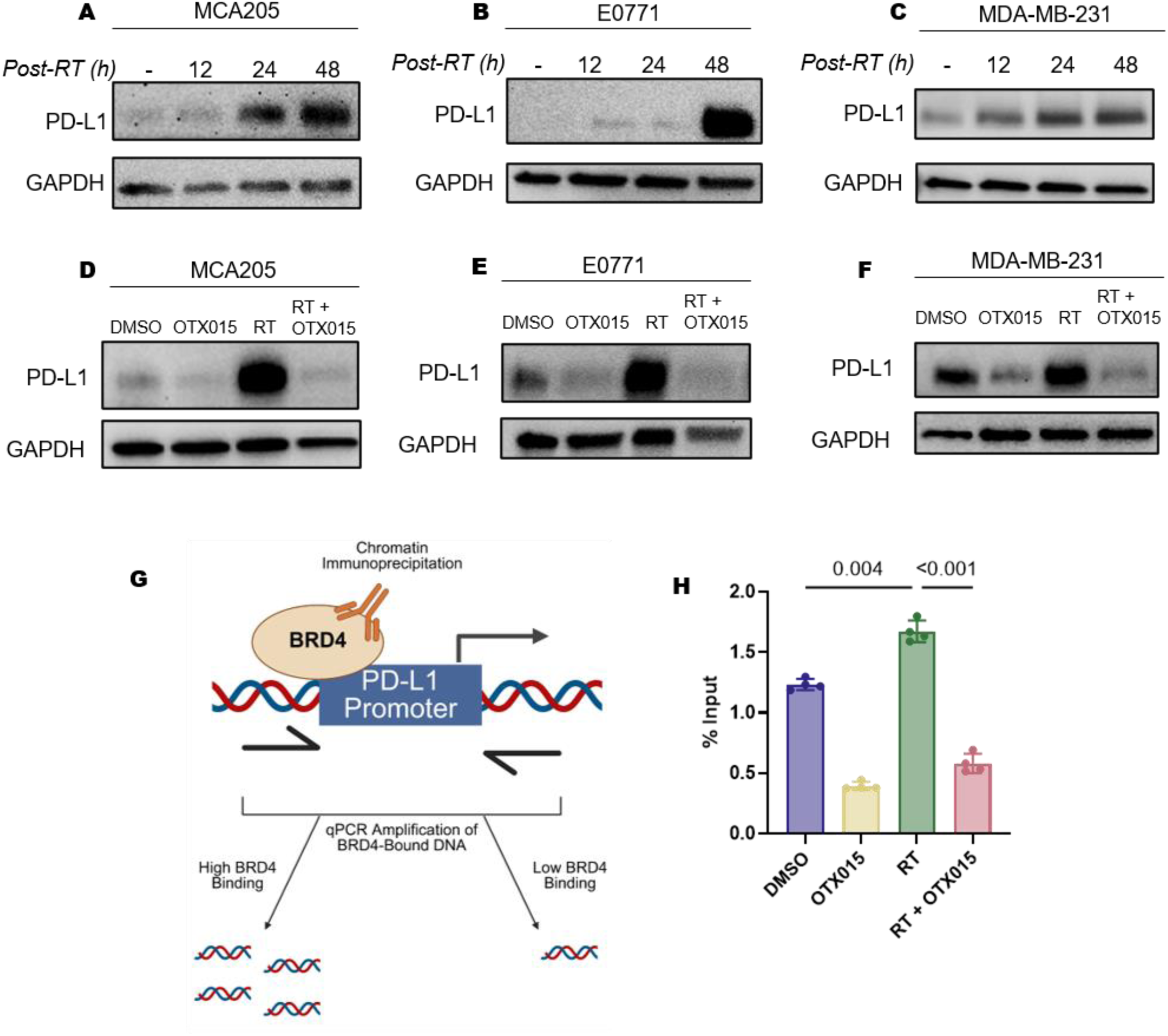
RT + OTX015 abrogates RT-Induced PD-L1 overexpression in tumor cells. (A-C): MCA205 (A), E0771 (B), and MDA-MB-231 (C) cells were treated with RT (8 Gy), cell lysates collected prior to RT, and at 12 hours, 24 hours and 48 hours post-RT, and subjected to Western blot analysis using anti-PD-L1 and GAPDH antibodies. (D-F): MCA205 (D), E0771 (E), and MDA-MB-231 (F) cells were treated with DMSO, OTX015 (0.5 µM), RT (8 Gy), or RT + OTX015, cell lysates were collected 48 hours post-treatment, and subjected to Western blot analysis with anti-PD-L1 and GAPDH. (G): Schematic representation of chromatin immunoprecipitation (ChIP) assay for quantifying BRD4 binding to the PD-L1 promoter. (H): MDA-MB-231 were cells treated with DMSO, OTX015 (0.5 µM), RT (8 Gy), and RT + OTX015 and subjected to ChIP assay using anti-BRD4 antibody followed by qPCR for PD-L1 promoter (n=4). Overall differences among groups were assessed using one-way ANOVA. Prespecified pairwise comparisons were performed using unpaired Welch’s *t*-tests to account for unequal variances.

## Discussion

The advent of immunotherapies such as ICIs has significantly improved survival in responding patients with many types of cancers (1–3). However, many patients with immunologically cold tumors such as BC and STS derive limited benefit from ICIs (4, 5). Even in patients with localized TNBC in whom immunotherapies are FDA-approved, the current SOC KEYNOTE-522 chemo-immunotherapy regimen requires prolonged co-administration of pembrolizumab with a multi-agent chemotherapy backbone, which results in high toxicity to patients. Thus, nearly 80% of patients receiving the KEYNOTE-522 regimen experience Grade 3 or higher toxicity with the chemotherapy backbone contributing to over 90% of the observed toxicity (54). Similarly, many patients with STS do not derive any benefit from existing immunotherapies and require prolonged administration of chemotherapy regimens that extract a high toxicity penalty with a modest benefit (5, 8, 9). Therefore, there is an urgent need to develop novel chemotherapy-free immunotherapy regimens which elicit a strong anti-tumor immune response with long term immunological memory against immunologically cold tumors such as BC and STS, thereby improving clinical outcomes in these patients while minimizing toxicity.

In this study, employing an *in vivo* drug screen, we identify that a short course of RT synergizes with BET inhibition and elicits a robust systemic anti-tumor immunity and long-term immunological memory across multiple immunologically-cold syngeneic mouse models, including BC and STS. Although RT alone and BET inhibition alone exhibit modest efficacy, the combination treatment achieves remarkable anti-tumor activity and long-term immunological memory in a tumor-specific fashion. The combination treatment orchestrates a favorable change in tumor microenvironment in both the local irradiated tumors and distant non-irradiated tumors as evidenced by increased infiltration of CD4+ and CD8+ T cells. The anti-tumor effects of RT + BET inhibition are mediated, at least in part, by cytotoxic T cells as depletion of CD8+ T cells abrogated tumor control afforded by the combination treatment.

The proposed novel combination has several advantages over existing treatment paradigms, including-

### Short treatment duration and low toxicity

Unlike existing systemic treatments for poorly immunogenic tumors such as BC and STS, which involve prolonged administration of chemotherapy-based regimen with or without immunotherapy, the proposed combination treatment involves less than a week of total treatment duration and does not require a chemotherapy backbone. Despite such short duration, the combination of RT + BET inhibition outperforms existing SOC anthracycline-based chemo-immunotherapy regimen without any evidence of toxicity in pre-clinical murine models. Since up to 90% of the toxicity from the existing SOC chemo-immunotherapy regimens is attributable to the chemotherapy backbone (54), a chemotherapy-free immunotherapy regimen is expected to dramatically reduce treatment-related toxicity in patients. Furthermore, administration of both RT and BET inhibitor at subtherapeutic doses contributes to the low overall toxicity of this regimen. The short treatment duration also has the potential to dramatically reduce financial toxicity and improve quality-of-life in patients by obviating the need for prolonged administration of high toxic chemotherapy-based regimens.

### Non-invasive route of administration

Since BET inhibitors are administered orally and radiation treatments are commonly delivered in a contact-free manner using a linear accelerator, the treatment is completely non-invasive. This has the potential to dramatically improve patient convenience and compliance.

### Activation of multiple steps involved in the generation of anti-tumor immunity

While immunotherapies such as ICIs have achieved remarkable clinical success, only a small subset of patients derive clinical benefit (55). This is likely due to the fact that the immune activation cycle is a complex process and involves multiple steps, including the release of tumor antigens, presentation of tumor antigens by APCs, priming and activation of cytotoxic T cells, trafficking and infiltration of T cell to the tumor microenvironment, recognition and killing of cancer cells and generation of memory T cells that mediate long term immunological memory (Figure 6). Since the existing clinically approved ICIs only impact a limited number of steps in this complex cycle, they only show benefit when the rate limiting step in the immune activation cycle in a given patient matches with the mechanism of action of the ICI. On the other hand, novel combination treatments that influence multiple steps in the immune activation cycle have the potential to induce a more robust anti-tumor immune response and benefit a larger proportion of patients.

**Figure 6:**
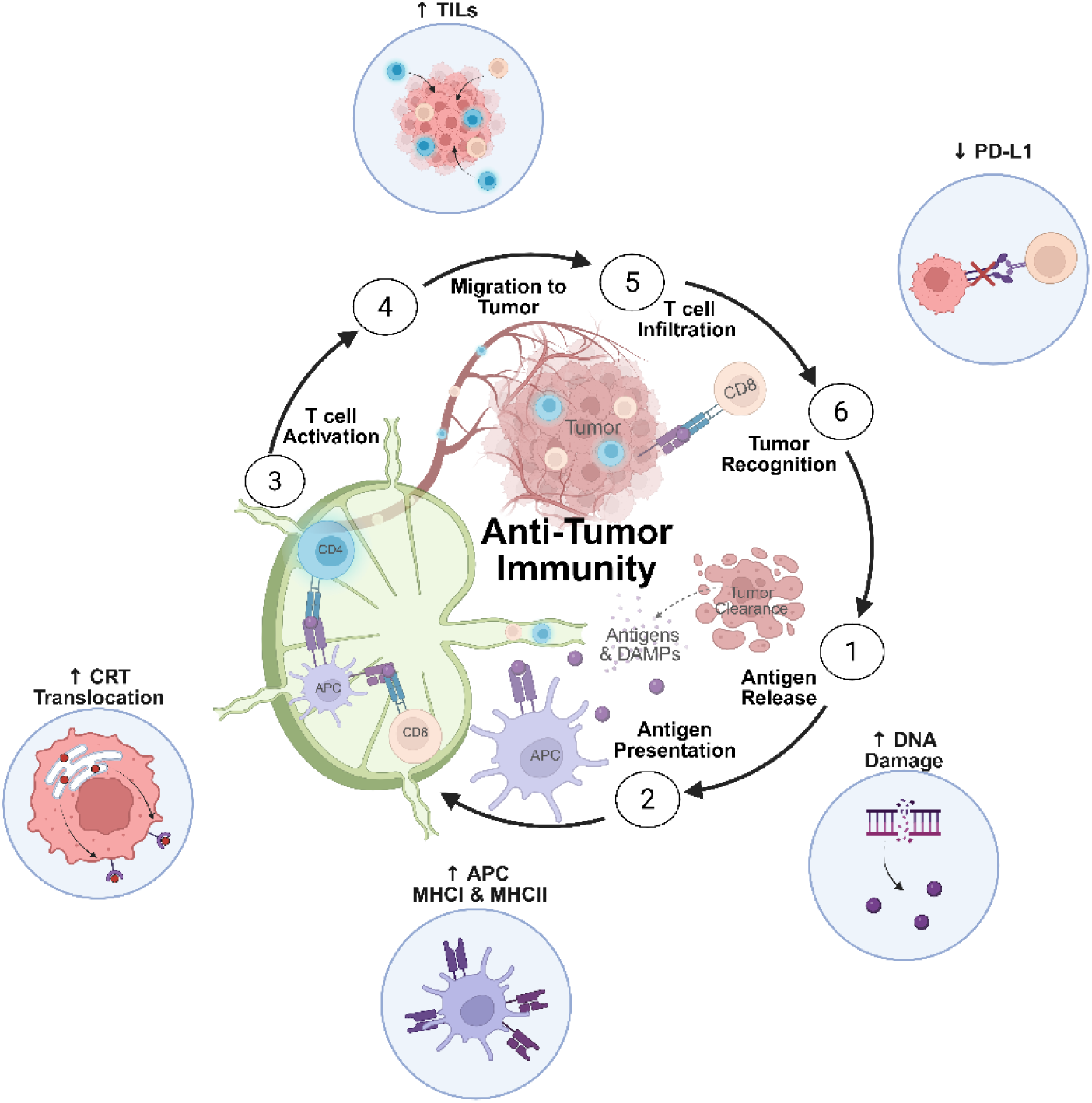
Schematic depiction of the immune activation cycle and the steps that are influenced by RT + OTX015. Development of anti-tumor immunity is a complex process with multiple steps that may serve as roadblocks against a productive immune response. RT + OTX015 influences multiple steps in this process to promote a robust anti-tumor immune response.

We have shown that RT + BET inhibition:

a. accentuates RT-induced DNA damage and release of micronuclei, which has the potential to activate the innate immune system through release of DAMPs,
b. attenuates repair of RT-induced DNA damage, which has the potential to generate neoantigens,
c. increases MHC class I and II expression on macrophages and translocation of calreticulin to plasma membrane, which promotes antigen presentation and T cell activation, and
d. abrogates RT-induced PD-L1 overexpression on tumor cells, which enhances T cell mediated cytotoxicity.

Recently, many pre-clinical studies have demonstrated that the combination of RT with other immunotherapies such as PD-L1 inhibition, CTLA4 inhibition, Flt3-Ligand and anti-amphiregulin improves both local and systemic control (19, 24, 56–63). Some of these studies have been successfully translated into clinical trials and have shown promising results (21, 64, 65). Our study, which shows that RT + BET inhibition positively influences several key steps involved in immune activation cycle, further advances this highly promising area of investigation.

### High translational potential

Since RT is routinely used as a common cancer treatment and several BET inhibitors are currently in clinical development with a well-established toxicity profile (66), our study has high potential for clinical translation. A major limitation of BET inhibitors is their systemic toxicity with prolonged administration, which hampered their clinical success and patient compliance (67). A major advantage of the proposed approach is that the total duration of BET inhibitor administration is less than 1 week. Furthermore, our approach relies on the ability of BET inhibition to accentuate the effects of RT on immune activation rather than on its own systemic effects. Consistent with this hypothesis, short course administration of BET inhibitors alone demonstrated minimal anti-tumor efficacy. This approach limits systemic toxicity of BET inhibition while maintaining high efficacy when combined with RT. This novel approach has the potential to lead to the revival of BET inhibitors in cancer therapy since the clinical development of many BET inhibitors has been abandoned after early phase human clinical trials due to the aforementioned limitations. Furthermore, the efficacy of the combination treatment against cancers of both epithelial and mesenchymal origin suggests broad applicability of this therapeutic approach against diverse cancers.

In addition, there is currently significant interest in employing metastases-directed therapies such as ablative radiation, in addition to the SOC systemic therapy, to prolong survival in patients with oligometastatic disease. However, two recent randomized control trials, NRG-BR002 and NRG-LU002, failed to show clinical benefit for metastases-directed therapy in patients with oligometastatic breast cancer and lung cancer, respectively (68, 69). However, since RT + BET inhibition stimulates systemic anti-tumor immunity and long term immunological memory (in addition to effectively controlling treated local tumors) in preclinical models, such an approach may have a higher potential for success than RT alone as metastases-directed therapy in oligometastatic cancer patients.

In summary, we describe a novel combination of RT + BET inhibition that stimulates robust and systemic anti-tumor immune response and long-term immunological memory in multiple and diverse pre-clinical murine cancer models by influencing multiple steps involved in the generation of anti-tumor immunity. Efforts are currently underway to translate these findings into a clinical trial.

## Methods

### Sex as a biological variable

Since all the murine models used in this study except MCA205 represent breast cancer models, and breast cancer is extremely uncommon in males, the study only employed female mice. Therefore, sex was not a biological variable in this study.

### Cell culture

4T1 (Cat #: CRL-2539), EMT6 (Cat #: CRL-2755), and E0771 (Cat #: CRL-341), and MDA-MB-231 (Cat#: HTB-26) were obtained from the American Type Culture Collection (ATCC). MCA205 was purchased from Millipore-Sigma (Cat #: SCC173). MXT+ cells were provided by Dr. Helmut Schonenberger. All cell lines were maintained in DMEM (Cat #: D6429, Sigma-Aldrich) with 10% FBS (Cat #: 35010CV, Corning) and 1% Penicillin-Streptomycin (Cat #: P0781, Sigma-Aldrich). Cells underwent regular mycoplasma testing using the MycoAlert® - Mycoplasma Detection Kit (Cat #: LT07-318, Lonza) to rule out mycoplasma contamination.

### Tumor Models

6-8 week old, female BALB/c (Inotiv), C57BL/6 (Charles River), or BDF1 (Charles River) mice were used for *in vivo* studies, as indicated. TNBC cell lines 4T1 (2 x 10^5^ cells), EMT6 (1 x 10^5^ cells), E0771 (5 x 10^5^ cells), and the ER+ cell line MXT+ (1 x 10^7^ cells) were suspended in 100 µL PBS and injected orthotopically in the mammary fat pad. The STS cell line MCA205 (5 x 10^5^ cells) was suspended in 100 µL PBS and injected subcutaneously in the flank. Tumors were measured twice weekly using digital calipers and tumor volumes calculated using the formula: volume = (large diameter x (small diameter)^2^) / 2.

For rechallenge studies, mice cured by RT + OTX015 were reinjected with the same tumor cell line in the contralateral site (mammary fat pad for BC models or flank for STS model) two months following initial tumor injection. Two months later, mice that resisted rechallenge were injected with a different syngeneic cell line, either 4T1 for mice cured of EMT6 or E0771 for mice cured of MCA205.

For immune depletion studies, mice were administered 200 µg anti-CD8 or anti-IgG depletion antibody intraperitoneally one day prior to tumor cell injection, the day of tumor cell injection, and twice weekly thereafter. Immune cell depletion was validated using flow cytometry on lymphocytes collected via submandibular cheek bleed.

### Irradiation

Tumor-directed x-ray radiation was administered using a small animal irradiator (X-RAD 320, Precision X-ray Inc) at 250 kV, 15 mA with a 2 mm aluminum filter at 1.39 Gy/min. Mice were placed under 1% to 5% isoflurane anesthesia during irradiation. 8 Gy irradiation was administered on days 2 and 4 or days 2, 4 and 6, as indicated.

For cells, x-ray irradiation was administered using the X-RAD 320 at 320 kV, 12.5 mA with a 2 mm aluminum filter at 3.64 Gy/min. A single fraction of 8 Gy was administered for *in vitro* studies.

### Western blotting

To assess protein expression, cultured cells were lysed using 1x RIPA buffer (Cat #: 20-188, Millipore) containing 1x protease inhibitor (Cat #: 11873580001, Roche) and 1x phosphatase inhibitor (Cat #: 04906837001, Roche). Total protein concentration was quantified using the Pierce™ BCA Protein Assay Kit (Cat #: 23225, Thermo Scientific). Denatured protein was run on SDS-PAGE at 80V and transferred onto a PVDF membrane. The membrane was then blocked in 5% milk in TBST, incubated with the indicated primary antibody overnight at 4°C. The membrane was then incubated with the indicated secondary antibody for 1 hour at room temperature. Proteins were detected using the SuperSignal™ West Femto Maximum Sensitivity Substrate (Cat #: 34096, Thermo Scientific).

### Immunofluorescence

Cells were seeded into 6-well plates containing coverslips. The following day, cells were treated with 0.5 µM OTX015 or DMSO. The following day, cells in the radiation arms received 8 Gy of RT as described above. Cells were harvested at the indicated timepoints, fixed with 4% paraformaldehyde for 10 minutes and washed three times with PBS. The cells were then blocked with 2 mg/mL BSA and 0.2% Triton-X in PBS for 30 minutes. Cells were washed three times with PBS and incubated with the primary antibody overnight at 4°C. The following day, the cells were washed three times with PBS and incubated with secondary antibody in 2 mg/mL BSA for 30 minutes at room temperature. Cells were washed three times with PBS, the coverslips were flipped onto labeled microscope slides with Vectashield Mounting Media with DAPI (Cat #: H-2000, Vector Laboratories Inc.). Slides were imaged on a Zeiss AxioImager M2 Microscope. Images were quantified using ImageJ.

For multiplexed immunofluorescent imaging of tissues, slide preparation, antibody staining, and image quantification were performed as previously described (70).

### Flow cytometry

Tumors were minced and digested in 2 mg/mL collagenase for 1 hour at 37°C. Cells were then passed through a 70 µM strainer and pelleted. RBCs were lysed with RBC lysis buffer for 10 minutes at room temperature and quenched with PBS. Cells in suspension (either from tumor digests or cultured cells) were washed with FACS buffer, blocked with Fc blocking antibody (Cat #: 553142, BD Pharmingen) in blocking buffer for 30 minutes on ice. Cells were then stained with fluorescently-conjugated antibodies for 1 hour at 4°C. For calreticulin staining, cells were stained with unconjugated primary antibody for 1 hour at 4°C followed by staining with a secondary fluorescent antibody for 30 minutes at 4°C. Cells were washed and analyzed on a Cytek SpectroFlo® Cytometer. Data was analyzed using FlowJo.

### Chromatin immunoprecipitation

MDA-MB-231 cells were treated with 0.5 µM OTX015 or DMSO. The following day, cells in the radiation arms received 8 Gy of RT as described previously. 6 hours later, cells were collected using the SimpleChIP® Enzymatic Chromatin IP Kit (Cat#: 9003, Cell Signaling Technologies). Cells were cross-linked, fragmented, and immunoprecipitated with BRD4 primary antibody according to the manufacturer instructions. QPCR using primers for the human PD-L1 promoter as previously described (53): forward 5’-GGGACCCTGAGCATTCTTAAA-3’, reverse 5’- GCCAACATCTGAACGCACC-3’.

### Statistics

Experiments were performed using biological replicates as indicated. All P-values are reported in Supplemental Table 3.

Tumor growth was analyzed using linear mixed-effects models with treatment and time included as fixed effects and mouse included as a random effect to account for repeated measurements. Pre-specified pairwise comparisons (e.g., treatments versus vehicle or versus combination therapy) were performed using estimated marginal means, with Benjamini-Hochberg correction applied within each comparison family. Because tumor volume variance increased with tumor size, analyses were additionally conducted on log-transformed tumor volumes to improve variance stability and model assumptions. P-values from the log-transformed models were selected for reporting, as these models demonstrated improved residual behavior and provide more reliable statistical inference in small-sample longitudinal tumor studies

For survival studies, overall survival differences among treatment groups were assessed using the long-rank test. Prespecified pairwise log-rank comparison were subsequently performed using defined reference groups, with p-values adjusted using Benjamini-Hochberg correction within each comparison family.

For all other experiments, statistics were calculated using GraphPad Prism 10. For experiments comparing two treatment conditions, significance was calculated using a 2-tailed student’s t-test with Welch’s correction to adjust for unequal variance. For experiments comparing more than two treatment conditions, overall differences among groups were assessed using a one-way ANOVA. Variance heterogeneity was evaluated using Brown-Forsyth test, and prespecified pairwise comparisons were performed using unpaired Welch’s t-tests when unequal variance were present.

## Study approval

All animal experiments were performed in accordance with UT Southwestern IACUC approved protocols.

## Data availability

All the primary data reported in this study can be accessed in the Supporting Data file.

## Author Contributions

PGA conceptualized this study. NM, CV, and PGA designed experiments and methodology. NM, CV, KP, SMNU conducted experiments and acquired data. NM, CV, YLL and PGA analyzed and interpreted the data. NM and PGA wrote the original manuscript. NM, CV, KP, SMNU, YLL and PGA reviewed and/or edited the final manuscript. PGA supervised this study.

## Acknowledgements

This work was funded by the Department of Defense Breast Cancer Research Program Breakthrough Awards (W81XWH2110112 and HT9425-25-1-0564) to PGA.

This study was supported by multiple Core Facilities at UT Southwestern Medical Center: (1) The Preclinical Imaging and Radiation Core Facility, supported by the Cancer Prevention & Research Institute of Texas under award S10 RP180770; (2) the Tissue Management Shared Resource, affiliated with the Simmons Comprehensive Cancer Center and supported in part by the National Cancer Institute award P30 CA142543; (3) the Metabolic Phenotyping Core; and (4) the Animal Resource Center (ARC). We thank Dr. Siyuan Zhang at UT Southwestern Medical Center for providing access to Cytek SpectroFlo® Cytometer, Zenith Epigenetics for providing ZEN-3694, Dr. Anthony Davis at UT Southwestern Medical Center for providing technical assistance with immunofluorescence experiments, and BioRender for graphics.

## Supplemental Material

**Figure S1:**
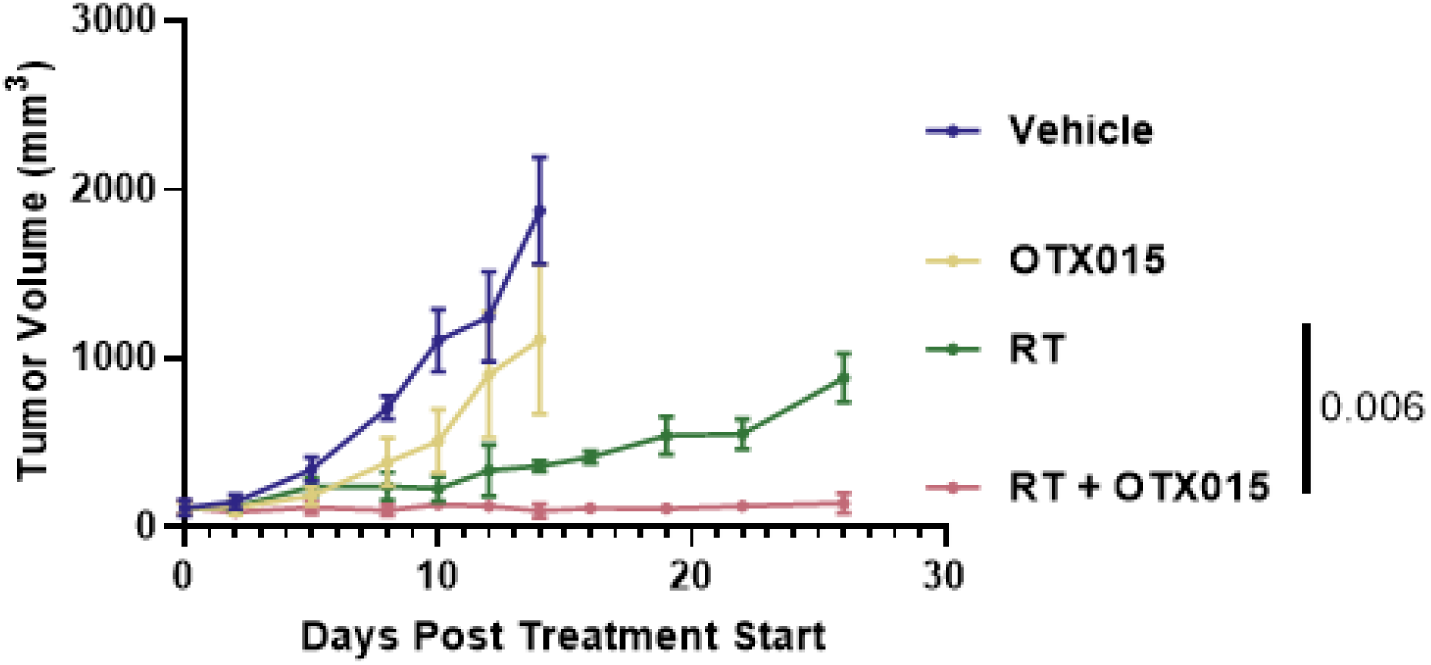
RT synergizes with OTX015 to inhibit the growth of MXT+ tumors. Female, BDF1 mice bearing ER+, MXT+ tumors were randomized when tumors reached 100-150 mm^3^ to the following treatment arms: Vehicle, OTX015 (100 mg/kg by oral gavage on days 1-6), RT (8 Gy on days 2, 4, and 6) or RT + OTX015. The results are plotted as average tumor volume for each treatment arm ± SD (n=5 mice/arm). P-values were determined using linear mixed-effects models, with Benjamini-Hochberg correction applied to prespecified pairwise comparisons.

**Figure S2:**
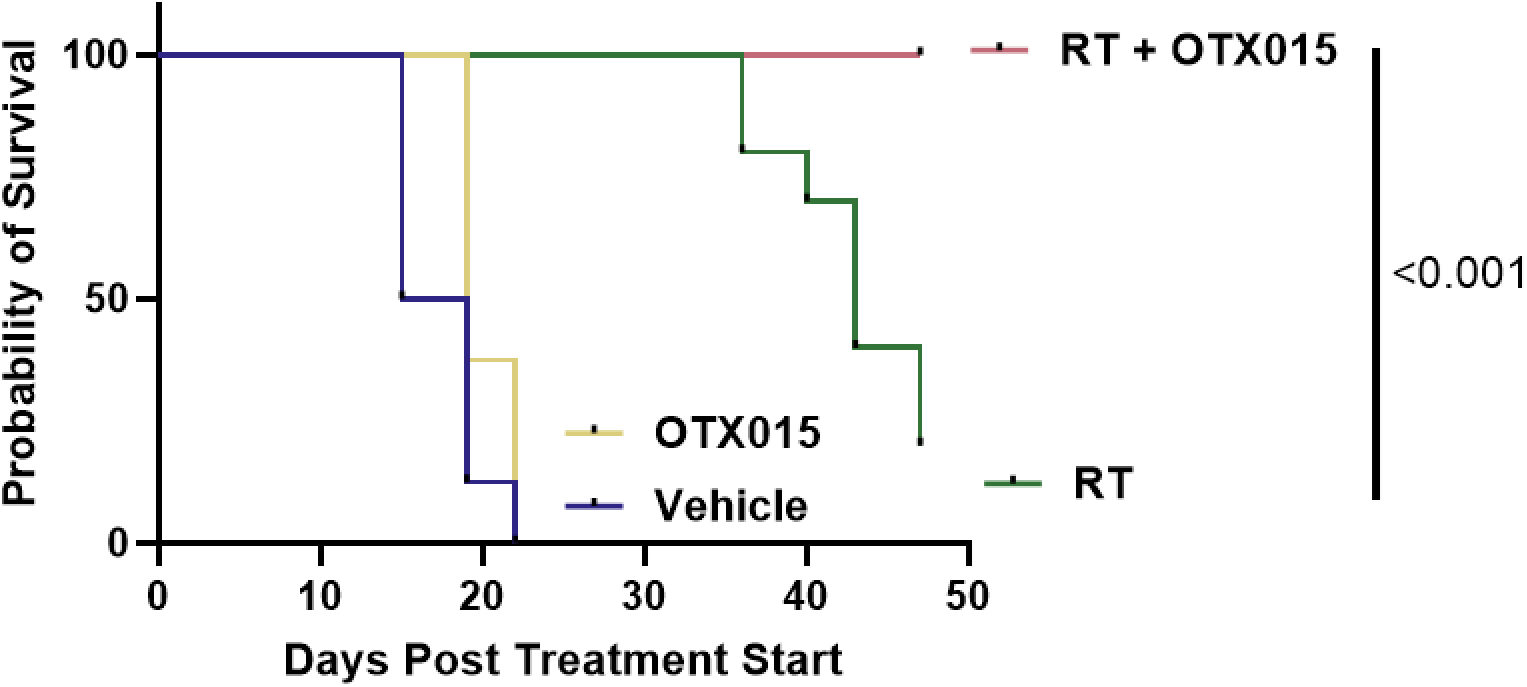
RT + OTX015 improves survival of MCA205 tumor bearing mice. Kaplan-Meier plot depicting survival of mice bearing MCA205 tumors treated with Vehicle, OTX015, RT, or RT + OTX015 as described in Figure 2D. At the end of treatment, 0/8 mice treated with the Vehicle, 0/8 mice treated with OTX015, 2/10 mice treated with RT and 10/10 mice treated with RT + OTX survived. P-values were derived from prespecified pairwise log-rank comparisons, with Benjamini-Hochberg correction applied for multiple comparisons.

**Figure S3:**
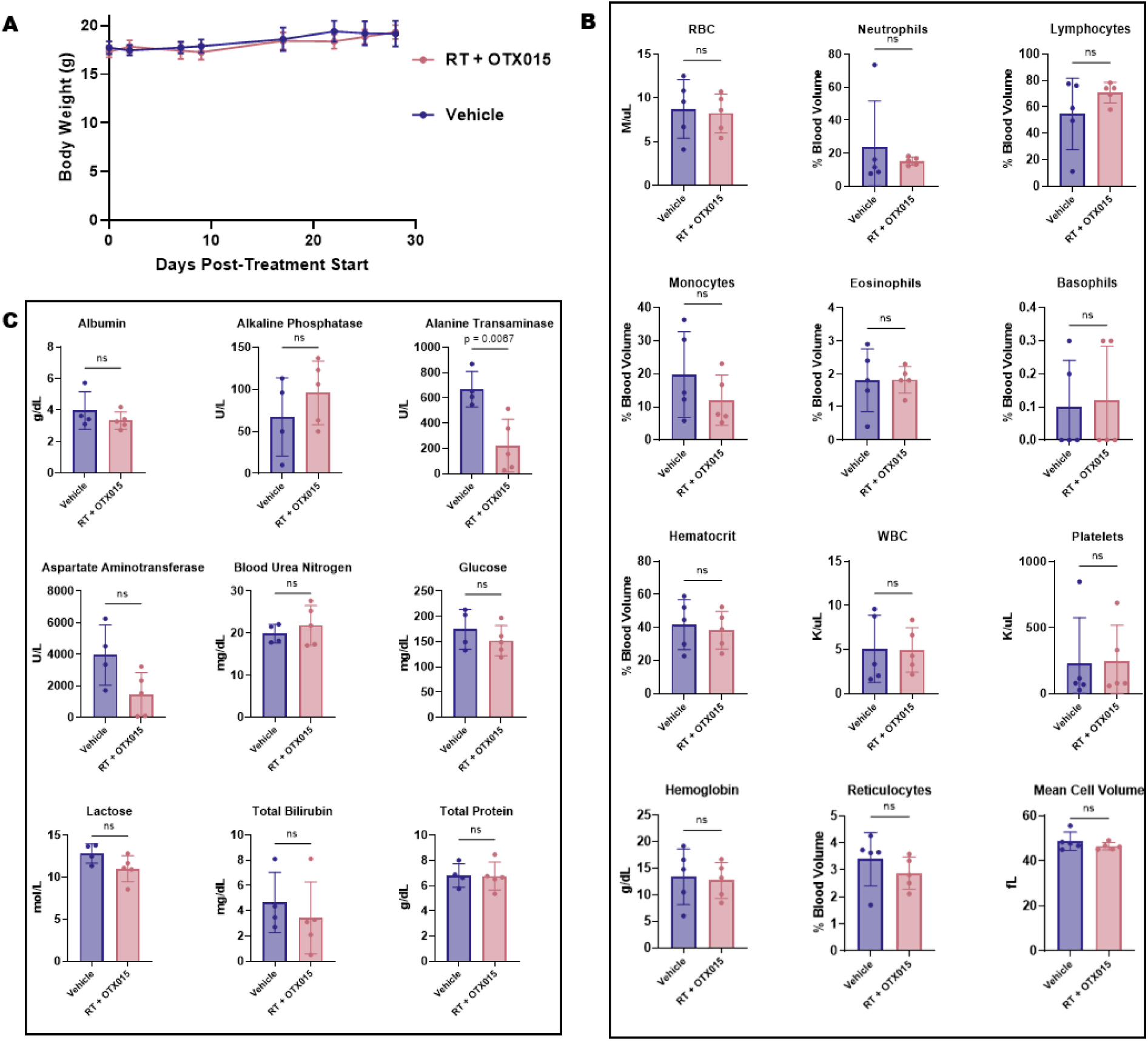
Mice treated with RT + OTX015 do not show any evidence of toxicity. Female BALB/c mice (n=5 mice/arm) without tumors were treated with Vehicle or RT (8 Gy on days 2 and 4 to the mammary fat pad) + OTX015 (100 mg/kg by oral gavage on days 1-4). Mice were monitored for 4 weeks and body weights (A), complete blood counts (B) and serum biochemistries (C) at the end of the study were plotted. P-values were determined using an unpaired, 2-tailed *t*-test with Welch’s correction.

**Figure S4:**
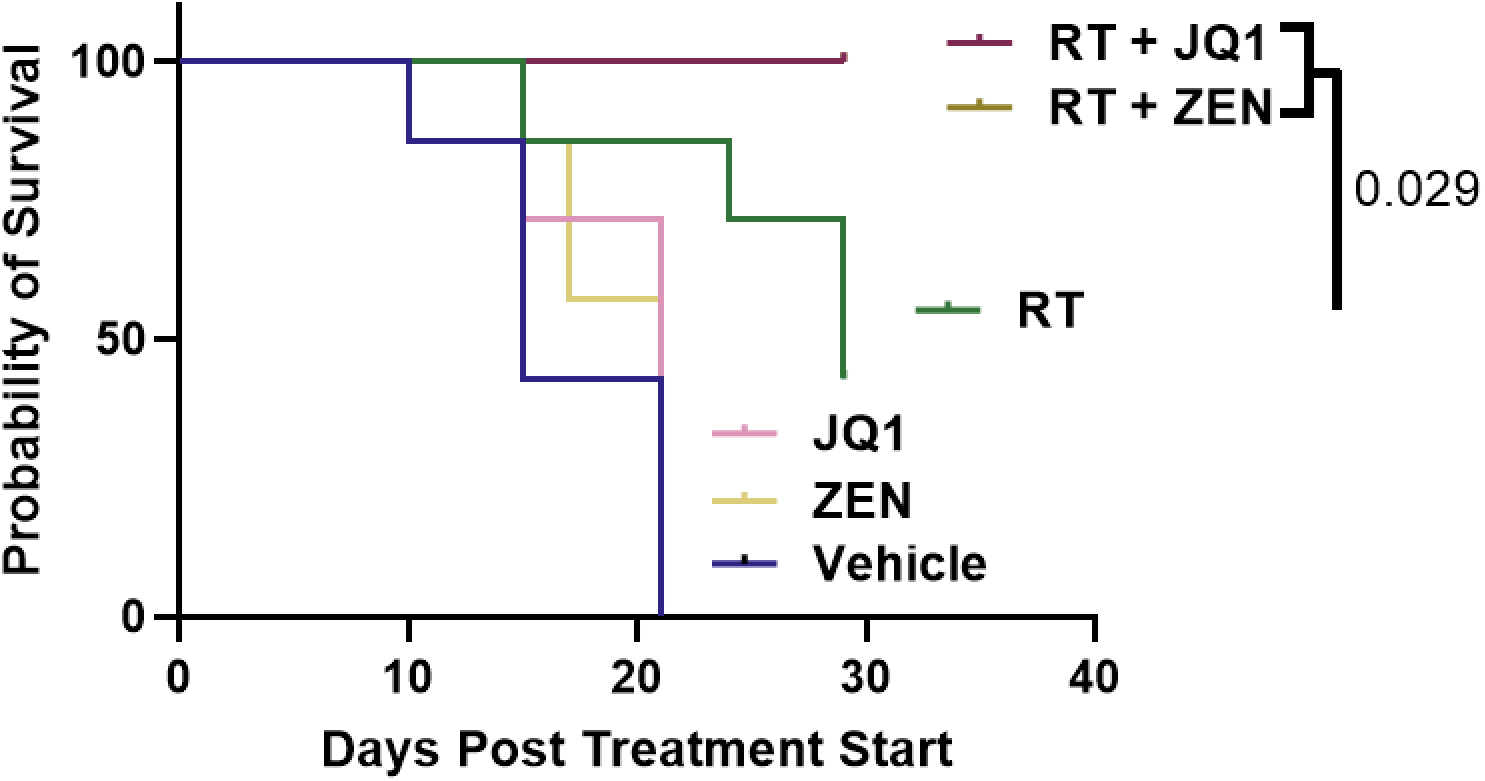
RT + ZEN-3694 and RT + JQ1 improve survival of MCA205 tumor bearing mice. Kaplan-Meier plot depicting survival of mice bearing MCA205 tumors treated with Vehicle, ZEN-3694, JQ1, RT, RT + ZEN-3694, or RT + JQ1 as shown in Figure 2E. At the end of treatment, 0/7 mice treated with vehicle, 0/7 mice treated with ZEN-3694, 0/7 mice treated with JQ1, 3/7 mice treated with RT, and 7/7 mice treated with RT + ZEN-3694 survived, and 7/7 mice treated with RT + JQ1 survived. P-values were derived from prespecified pairwise log-rank comparisons, with Benjamini-Hochberg correction applied for multiple comparisons.

**Supplemental Table 1:**
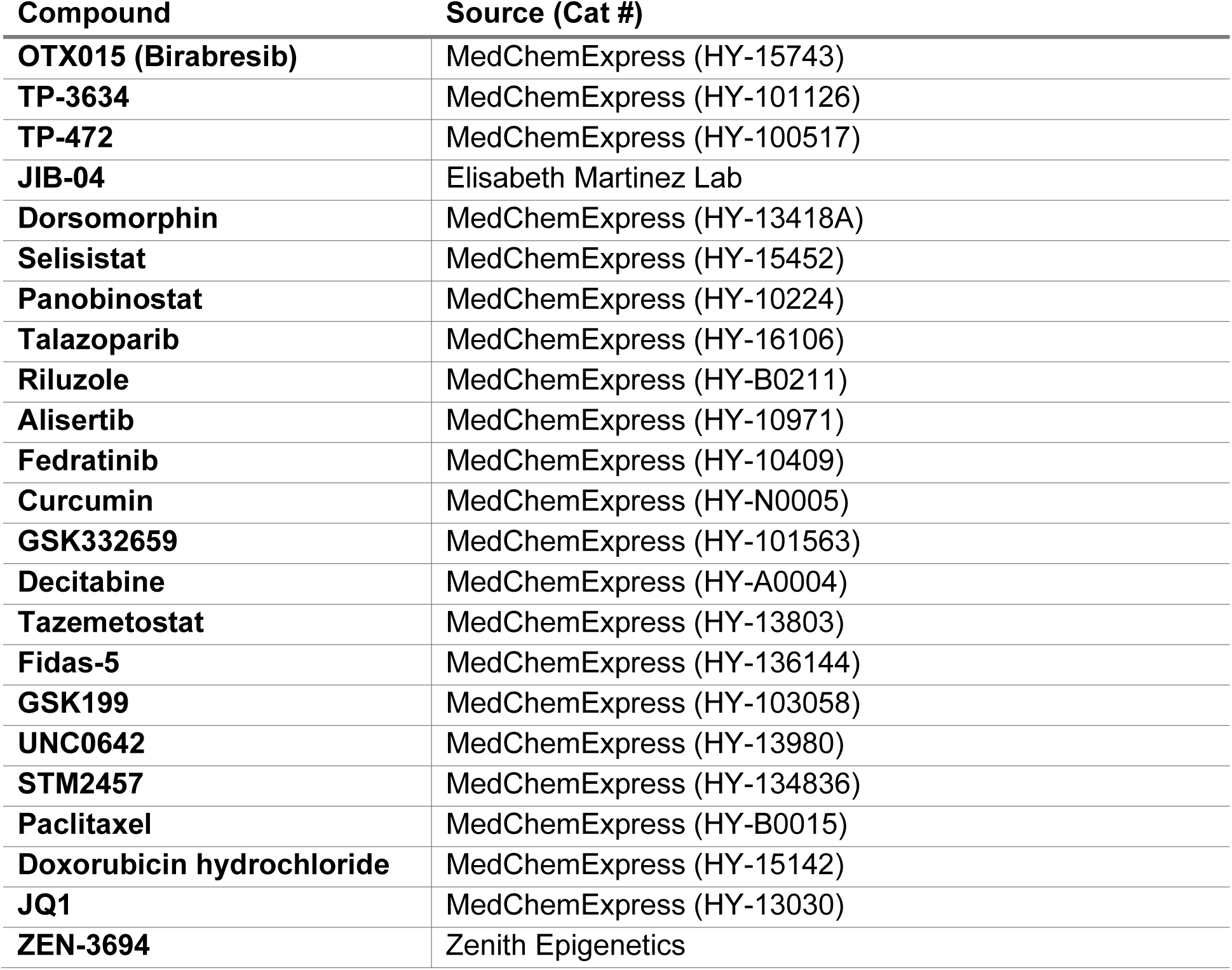
Chemicals and Drugs.

**Supplemental Table 2:** Antibodies.

| Target | Antibody | Dose/Dilution | Application | Source (Cat #) |
| --- | --- | --- | --- | --- |
| **Therapeutic Antibodies** |  |  |  |  |
| **PD1** |  | 200 µg | *In vivo* therapeutic | BioXCell (BE0146) |
| **IgG** |  | 200 µg | *In vivo* depletion | BioXCell (BE0089) |
| **CD8** |  | 200 µg | *In vivo* depletion | BioXCell (BE0004) |
| **Primary Antibodies** |  |  |  |  |
| **PDL1** | Rabbit anti-mouse PDL1, clone EPR20529 | 1:1,000 | Western blot | Abcam (ab213480) |
| **PDL1** | Rabbit anti-human PDL1, clone E1L3N | 1:1,000 | Western blot | Cell Signaling Technologies (13684) |
| **GAPDH** | Rabbit anti-GAPDH, clone 14C10 | 1:1,000 | Western blot | Cell Signaling Technologies (2118) |
| **BRD4** | Rabbit anti-BRD4, clone E1Y1P | 1:100 | Chromatin Immunoprecipitation | Cell Signaling Technologies (83375) |
| **Calreticulin** | Rabbit anti-Calreticulin, clone D3E6 | 1:100/<br>1:1,000 | Flow cytometry/<br>Immunofluorescence | Cell Signaling Technologies (12238) |
| **CD45** | Rat anti-mouse CD45, clone 30-F11; PE-Cy7 | 1:50 | Flow cytometry | BD Biosciences (552848) |
| **CD11b** | Rat anti-CD11b, clone M1/70; APC-Cy<sup>TM</sup>7 | 1:50 | Flow cytometry | BD Biosciences (557657) |
| **F480** | Rat anti-mouse F480, clone BM8; BV605 | 1:50 | Flow cytometry | BioLegend (123133) |
| **MHCI** | Rat anti-mouse H-2Kb, clone AF6-88.5; Pacific Blue | 1:50 | Flow cytometry | BioLegend (116514) |
| **MHCII** | Rat anti-mouse I-A/I-E, clone 2G9; FITC | 1:50 | Flow cytometry | BD Biosciences (562009) |
| **γH2AX** | Mouse anti-phospho-Histone H2A.X (Ser139), clone JBW301; unconjugated | 1:1,000 | Immunofluorescence | Sigma-Aldrich (05-636) |
| **CD4** | Rabbit anti-CD4,<br>clone EPR19514;<br>unconjugated | 1:8,000 | Immunofluorescence | Abcam<br>(ab183685) |
| **CD8** | Rabbit anti-CD8α, clone<br>D4W2Z;<br>unconjugated | 1:8,000 | Immunofluorescence | Cell Signaling<br>Technologies<br>(98941) |
| **FoxP3** | Rabbit anti-FoxP3, clone<br>1054C | 1:2,000 | Immunofluorescence | R&D<br>(MAB8214) |
| **Secondary Antibodies** |  |  |  |  |
| **Mouse IgG** | Horse anti-Mouse IgG; HRP-linked | 1:10,000 | Western Blot | Cell Signaling<br>Technologies<br>(7076) |
| **Rabbit IgG** | Goat anti-rabbit<br>IgG; HRP-linked | 1:10,000 | Western Blot | Cell Signaling<br>Technologies<br>(7074) |
| **Mouse IgG** | Donkey anti-mouse IgG;<br>Alexa Fluor® 488 | 2 drops/mL | Immunofluorescence | Invitrogen<br>(R37114) |
| **Rabbit IgG** | Donkey anti-rabbit IgG; Alexa<br>Fluor® 594 | 2 drops/mL | Immunofluorescence | Invitrogen<br>(R37119) |

**Supplemental Table 3:** Statistics.

| Figure | Comparison Group | Reference Group | Adjusted p-value |
| --- | --- | --- | --- |
| **1E** | OTX015 | Vehicle | 0.873 |
|  | PD1 + PTX + DOX | Vehicle | 0.135 |
|  | RT | Vehicle | 0.013 |
|  | RT + OTX015 | Vehicle | <0.001 |
|  | OTX015 | RT + OTX015 | <0.001 |
|  | PD1 + PTX + DOX | RT + OTX015 | <0.001 |
|  | RT | RT + OTX015 | <0.001 |
|  | Vehicle | RT + OTX015 | <0.001 |
| **1G** | PD1 + OTX015 | RT + OTX015 | <0.001 |
| **2C** | OTX015 | Vehicle | 0.327 |
|  | RT | Vehicle | 0.088 |
|  | RT + OTX015 | Vehicle | <0.001 |
|  | OTX015 | RT + OTX015 | <0.001 |
|  | RT | RT + OTX015 | <0.001 |
|  | Vehicle | RT + OTX015 | <0.001 |
| **2D** | OTX015 | Vehicle | 0.764 |
|  | RT | Vehicle | 0.384 |
|  | RT + OTX015 | Vehicle | 0.001 |
|  | OTX015 | RT + OTX015 | 0.001 |
|  | RT | RT + OTX015 | 0.004 |
|  | Vehicle | RT + OTX015 | 0.001 |
| **2E** | JQ1 | Vehicle | 0.043 |
|  | RT | Vehicle | 0.027 |
|  | RT + JQ1 | Vehicle | <0.001 |
|  | RT + ZEN | Vehicle | <0.001 |
|  | ZEN | Vehicle | 0.023 |
|  | JQ1 | RT + ZEN | <0.001 |
|  | RT | RT + ZEN | <0.001 |
|  | RT + JQ1 | RT + ZEN | 0.783 |
|  | Vehicle | RT + ZEN | <0.001 |
|  | ZEN | RT + ZEN | <0.001 |
|  | JQ1 | RT + JQ1 | <0.001 |
|  | RT | RT + JQ1 | <0.001 |
|  | RT + ZEN | RT + JQ1 | 0.784 |
|  | Vehicle | RT + JQ1 | <0.001 |
|  | ZEN | RT + JQ1 | <0.001 |
| **3B** | RT | RT + OTX015 | 0.030 |
| **3C** | RT | RT + OTX015 | 0.006 |
| **3E** | RT | RT + OTX015 | 0.038 |
| **3F** | RT | RT + OTX015 | <0.001 |
| **3G** | RT | RT + OTX015 | 0.024 |
| **3H** | RT | RT + OTX015 | 0.006 |
| **3I** | CD8 Depletion | IgG Depletion | 0.008 |
| **3J** | CD8 Depletion | IgG Depletion | 0.001 |
| **3K** | CD8 Depletion | IgG Depletion | <0.001 |
| **4B** | DMSO | RT + OTX – 30 min | <0.001 |
|  | OTX015 | RT + OTX – 30 min | <0.001 |
|  | RT – 30 min | RT + OTX – 30 min | 0.004 |
|  | RT + OTX015 – 120 min | RT + OTX – 30 min | 0.072 |
|  | DMSO | RT + OTX015 – 120 min | <0.001 |
|  | OTX015 | RT + OTX015 – 120 min | <0.001 |
|  | RT – 120 min | RT + OTX015 – 120 min | 0.004 |
| **4D** | DMSO | RT + OTX015 | <0.001 |
|  | OTX015 | RT + OTX015 | <0.001 |
|  | RT | RT + OTX015 | <0.001 |
| **4E** | Vehicle | RT + OTX015 | <0.001 |
|  | OTX015 | RT + OTX015 | <0.001 |
|  | RT | RT + OTX015 | 0.046 |
| **4F** | Vehicle | RT + OTX015 | 0.002 |
|  | OTX015 | RT + OTX015 | <0.001 |
|  | RT | RT + OTX015 | 0.002 |
| **4H** | DMSO | RT + OTX015 | <0.001 |
|  | OTX015 | RT + OTX015 | <0.001 |
|  | RT | RT + OTX015 | <0.001 |
| **4I** | DMSO | RT + OTX015 | <0.001 |
|  | OTX015 | RT + OTX015 | <0.001 |
|  | RT | RT + OTX015 | <0.001 |
| **5H** | DMSO | RT | 0.004 |
|  | OTX015 | RT | <0.001 |
|  | RT + OTX015 | RT | <0.001 |
|  | DMSO | RT + OTX015 | <0.001 |
|  | OTX015 | RT + OTX015 | 0.047 |
| **S1** | OTX015 | Vehicle | <0.001 |
|  | RT | Vehicle | <0.001 |
|  | RT + OTX015 | Vehicle | <0.001 |
|  | OTX015 | RT + OTX015 | <0.001 |
|  | RT | RT + OTX015 | 0.006 |
|  | Vehicle | RT + OTX015 | <0.001 |
| **S2** | OTX015 | Vehicle | 0.052 |
|  | RT | Vehicle | <0.001 |
|  | RT + OTX015 | Vehicle | <0.001 |
|  | OTX015 | RT + OTX015 | <0.001 |
|  | RT | RT + OTX015 | <0.001 |
|  | Vehicle | RT + OTX015 | <0.001 |
| **S4** | JQ1 | Vehicle | 0.314 |
|  | RT | Vehicle | 0.005 |
|  | RT + JQ1 | Vehicle | 0.001 |
|  | RT + ZEN | Vehicle | 0.001 |
|  | ZEN | Vehicle | 0.401 |
|  | JQ1 | RT + ZEN | 0.001 |
|  | RT | RT + ZEN | 0.029 |
|  | RT + JQ1 | RT + ZEN | >0.999 |
|  | Vehicle | RT + ZEN | 0.001 |
|  | ZEN | RT + ZEN | 0.001 |
|  | JQ1 | RT + JQ1 | 0.001 |
|  | RT | RT + JQ1 | 0.029 |
|  | RT + ZEN | RT + JQ1 | >0.999 |
|  | Vehicle | RT + JQ1 | 0.001 |
|  | ZEN | RT + JQ1 | 0.001 |

## Notes

### Competing Interest Statement

The authors have declared no competing interest.

